# *O*^6^-Alkylguanine-DNA Alkyltransferase Maintains Genomic Integrity During Peroxynitrite-Mediated DNA Damage by Forming DNA-Protein Crosslinks

**DOI:** 10.1101/2024.02.22.581550

**Authors:** Shayantani Chakraborty, Gargi Mukherjee, Anindita Chakrabarty, Goutam Chowdhury

**Affiliations:** Department of Life Science, Shiv Nadar Institution of Eminence (DTU), Greater Noida, UP, India; Independent Researcher, Greater Noida, UP, India

**Keywords:** Inflammation, Peroxynitrite, AGT, MGMT, DNA-protein crosslink

## Abstract

Inflammation is an early immune response against invading pathogens and damaged tissue. Although beneficial, uncontrolled inflammation leads to various diseases and may be fatal. Peroxynitrite (PN) is a major reactive nitrogen species (RNS) generated during inflammation. It produces various DNA lesions including labile 8-nitroguanine which spontaneously converts into abasic sites resulting in DNA strand breakage. Here, we report the discovery of a previously unrecognized function of the human repair protein *O*^6^-alkylguanine-DNA alkyltransferase (hAGT or MGMT). We showed that hAGT through its active site nucleophilic Cys145 thiolate can spontaneously react with 8-nitroguanine in DNA to form a stable DNA-protein crosslink (DPC). Interestingly, the process of DPC formation provides protection from PN-mediated genome instability. The Cys145-mutant of hAGT failed to form DPC and provide protection against inflammation-associated, PN-mediated cytotoxicity. Gel shift, dot blot and UV-Vis assays showed formation of a covalent linkage between PN-damaged DNA and hAGT through its active site Cys145. Finally, expression of hAGT was found to be significantly increased by induced macrophages and PN. The data presented here clearly demonstrated hAGT as a dual function protein that along with DNA repair is capable of maintaining genomic integrity and providing protection from the toxicity caused by PN-mediated DNA damage. Although DPCs may seem detrimental, there are multiple systems in place in normal cells for their repair.

## Introduction

Living beings have developed the immune system as part of their survival strategy. There are two complementary immune systems in vertebrates, innate and adaptive.[1, 2] The innate immune system elicits an immediate non-selective response in the form of inflammation against invading pathogens, damaged tissues and foreign agents. Although beneficial, uncontrolled inflammation may cause many diseases including diabetes, cancer, organ damage/failure and mortality. During the recent COVID-19 pandemic, it led to severe lung injury, hospitalization and fatality.[3–9]

Inflammation is associated with release of reactive oxygen (ROS) and nitrogen species (RNS) from invading neutrophils and macrophages as well as target cells.[10, 11] These are not only detrimental for pathogen-infested target cells but also for surrounding host cells, causing damage to cellular macromolecules including DNA, proteins and lipids. Peroxynitrite (PN) is a biological RNS formed in significant amounts at sites of inflammation by the rapid diffusion-controlled combination of superoxide (O ^·-^) and nitric oxide (^•^NO) radicals.[12–16] Apart from protein nitrosylation, PN also induces DNA damage.[17, 18] Various DNA lesions formed due to PN exposure include 8-nitroguanine (8-NO_2_-G), 8-oxo-7,8-dehydroguanine (8-oxo-G), 5-guanidino-4-nitroimidazole (NIm) and GT cross-links.[19–21] The major lesion 8-NO_2_-G is labile (half-life ∼1.6 and 2.4 h for single- and double-stranded DNA, respectively) and undergoes spontaneous depurination to form abasic sites which ultimately lead to DNA strand breaks.[22] 8-NO_2_-G is also known to be mutagenic.[23]

*O*^6^-alkylguanine-DNA alkyltransferase (AGT, MGMT) is a ubiquitous repair protein involved in the direct reversal of mutagenic DNA lesion *O*^6^-methylguanine and to a lesser extent *O*^4^-methylthymine to the natural base guanine and thymine, respectively.[24, 25] AGT contains an active site reactive cysteine (Cys145), in which the pK_a_ of thiol group is ∼5 owing to the presence of a catalytic triad.[26, 27] Upon binding to the *O*^6^-alkylguanine lesion in DNA, the alkyl group is stoichiometrically transferred to the Cys145 thiolate following an SN_2_ type mechanism. Alkylating agents induce AGT expression as a cytoprotective mechanism.[28] AGT is a well-recognized predictive biomarker of resistance to alkylating chemotherapeutic agents.[29] AGT can also form DNA-protein crosslinks (DPCs) with abasic sites and in the presence of cisplatin.[30–32] Although DPCs are bulky adducts and initially thought to cause replication blocks, there are DNA-stimulated proteases capable of converting these into DNA-peptide crosslinks that can be repaired or bypassed by DNA polymerases and may or may not miscode.[33, 34] Hence, DPCs are not necessarily always cytotoxic or mutagenic.

We have recently shown that the cytotoxicity resulting from PN-mediated DNA damage is mitigated by reduced glutathione (GSH).[21] The nitro group of the labile 8-NO_2_-G lesion formed in DNA following PN exposure is substituted by the sulfhydryl group of GSH resulting in a stable GS-G adduct.[21] Because the pK_a_ of the thiol group in GSH is 8.7,[35] and GSH has no DNA binding affinity, we hypothesized that DNA-binding proteins having reactive Cys with lower pK_a_ in their active sites may provide similar or enhanced protection. The repair protein AGT, having an active site Cys with thiol pK_a_ of ∼5,[36] strong affinity for DNA and ability to react with its substrate in stoichiometric amounts, seems like a lucrative candidate.[24] It has been reported that Cys145 of AGT undergoes reversible S-nitrosylation in presence of NO resulting in its deactivation and fast removal following ubiquitination.[37, 38] This intrigued us even more to see if AGT can maintain genomic integrity by providing protection from PN-mediated DNA strand breakage despite having an opposing deactivation and removal pathway during inflammation. Herein, using various biochemical and cellular assays we confirmed that hAGT (human AGT) indeed provides protection from the cytotoxicity caused by PN-mediated DNA damage by forming a DNA-AGT crosslink.

## Results and discussion

### hAGT protects mammalian cells from genomic integrity, cytotoxicity, and other stress responses such as senescence and polyploidy triggered by activated peripheral blood mononuclear cells (PBMC) or induced macrophages

To determine if hAGT can protect target mammalian cells from the cytotoxic effects of reactive species generated during inflammation, we used two cell culture-based model systems. In model system 1 differentially labelled (with fluorescent CellTracker^TM^ dyes) target cells and activated PBMC or induced macrophages were co-cultured. Target cells were labelled with blue dye, induced macrophages with orange dye and activated PBMC with green dye for visualization and tracking. In the model system 2, cells were grown in transwell chambers, with target cells in the outer chamber and activated PBMC or induced macrophages in the inner chamber. Activated PBMC or induced macrophages triggers generation of ROS and RNS leading to DNA damage (causing loss of genomic integrity) and cytotoxicity in target cells.[39] The co-culture model system 1 allows these released reactive species from activated PBMC or induced macrophages to immediately act upon the neighbouring target cells. In the transwell model system 2 the reactive species produced by activated immune cells in the inner chamber diffuse through the transwell membrane and act upon the target cells grown in the outer chamber. We measured the cytotoxic effects of these reactive species on target cells by cell viability and apoptosis assays. We performed single cell gel electrophoresis/comet assay to measure the extent of DNA damage and examined its effect on cell cycle by detection of growth-arrested/senescent cells and genomic instability by estimating polyploid cells. Formation of reactive oxidants from activated immune cells and target cells were measured by mitochondrial superoxide formation with “MitoSOX” reagent, a generic indicator of cellular oxidative stress.[40] Although these models do not completely mimic *in vivo* inflammation, they are practical and informative alternatives to available animal models for measuring toxicological outcomes.

In the co-culture model system 1, activated PBMC showed significant reactive species production within 1 h of activation/induction that persisted even after 6 h (Figure S1). One interesting observation was the detection of MitoSOX signal in the target MDA-MB-231 cells after 6 h of co-culture with PBMC, possibly through generation of NO and superoxide in target cells. The MDA-MB-231 cell was chosen because it has undetectable level of intrinsic AGT.[41] When transfected with empty pEGFP-N1 plasmid and co-cultured with activated PBMC for 0, 12, 36 h (Figure 1A and Figure S2), we observed significant reduction in cell number compared to the non-activated PBMC control. Interestingly, in case of hAGT-pEGFP-N1-transfected MDA-MB-231 cells overexpressing hAGT (Figure S3), the cytotoxic effect of the activated PBMC was reversed as the cell number was comparable to the non-activated control and greater than the empty vector control (Figure 1A and Figure S2). Such “protective effect” from PBMC-mediated cytotoxicity was abrogated by the addition of an inhibitor of hAGT, *O*^6^-benzylguanine to the AGT-overexpressing MDA-MB-231 cells (Figure 1A and Figure S2).[42] Instead of wild type hAGT, when a Cys145-mutated hAGT (C145S-hAGT-pEGFP-N1) was overexpressed in MDA-MB-231 cells (Figure S3), we were unable to protect the cells from the cytotoxic effect of the activated PBMC (Figure 1A and Figure S2). Using M1-induced THP-1 macrophages (Figure S4) instead of PBMC yielded similar results (Figure 1B and Figure S5).

**Figure 1.**
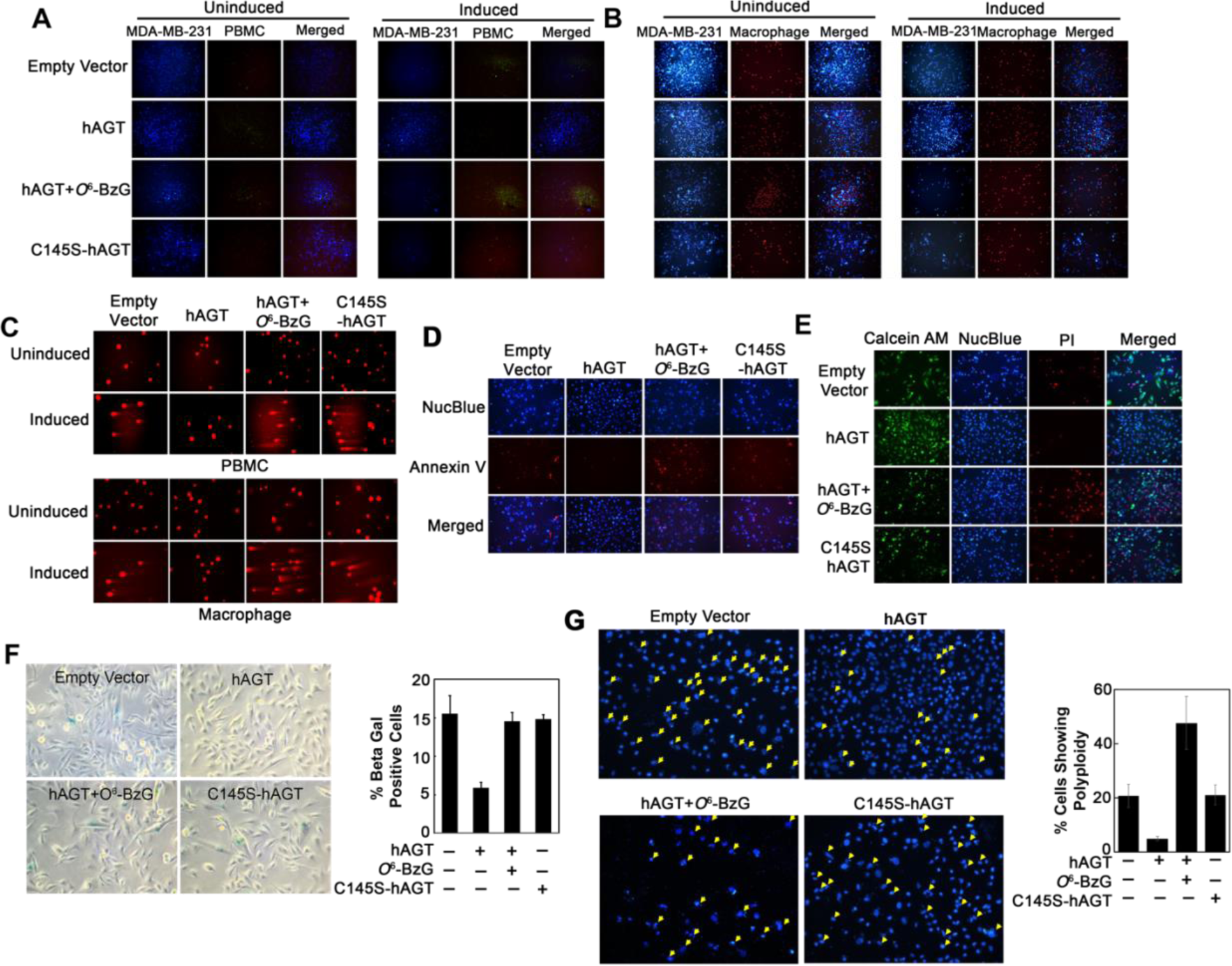
Effect of hAGT on activated PBMC- or induced macrophage-mediated DNA damage, cytotoxicity and stress response. A) Live cell images of co-cultured MDA-MB-231 and activated PBMC cells that were labelled with cell tracker blue and green dyes, respectively. MDA-MB-231 cells were transfected with empty pEGFPN-1, hAGT pEGFPN-1 or C145S-mutated hAGT pEGFPN-1 vectors for overexpression of the corresponding proteins and subsequently co-cultured for 12 h with activated PBMC. In one set of experiments, hAGT pEGFPN-1-transfected MDA-MB-231 cells were also treated with the known inhibitor of AGT *O*^6^-BzG 6 h post transfection. B) Same as in A with the exception of M1 polarized macrophages were used instead of PBMC. C) MDA-MB-231 cells transfected with empty pEGFPN-1, hAGT pEGFPN-1 or C145S-hAGT pEGFPN-1 constructs were cultured in transwell plates in presence of activated PBMC or induced M1 macrophages for 12 h and analysed by comet assay. The comet tail length indicates extent of DNA damage. D) Same as in A with the exception that instead of co-culture with PBMC, MDA-MB-231 cells were treated with PN (2.5 μM) 24 h post transfection for 4 min on ice. 48 h after PN treatment cells were imaged with NucBlue (nuclear staining of live cells) or Annexin V dye (apoptotic cells). E) Same as in D with the exception that imaging was done after adding Calcein-AM (live cells), NucBlue (nuclear staining), and PI (dead cells). F) SA-β-galactosidase immunohistochemistry was performed after 48 h of PN-treatment with MDA-MB-231 cells expressing different hAGT constructs as mentioned above. G) Hoechst staining was done with PN-treated MDA-MB-231 cells expressing different hAGT constructs as mentioned followed by fluorescence microscopy. The arrowheads indicate polyploid nuclei. Images were acquired at 10X magnifications for A and B and 20X for the rest. Quantitation of fluorescence signals was done with ImageJ software (National Institute of Health, Bethesda, MD, USA). Each bar graph is an average of three replicates with SE.

To determine if the cytotoxic effect of activated PBMC is associated with DNA damage and loss of genomic integrity in target cells, comet assay was performed with the MDA-MB-231 cells from the transwell model system 2. As expected, within 6 h of exposure of activated macrophages and PBMC, comet formation was observed in the target cells, possibly due to the release of diffusible factors such as ROS, RNS and cytokines which can travel through the transwell membrane and impact target cells (Figure 1C).

In order to directly correlate these cytoprotective functions of hAGT with PN-mediated damages, we repeated the above set of experiments with PN instead of activated immune cells. We treated hAGT-overexpressing MDA-MB-231 cells with 2.5 μM PN for 4 min on ice and cultured for another 48 h before imaging with various dyes. PN treatment resulted in apoptosis (Annexin V-positive cells) of empty vector-transfected MDA-MB-231 cells (Figure 1D). There was notable protection from apoptosis upon wild type hAGT overexpression, as indicated by reduced number of Annexin V-positive apoptotic cells and increased number of live cells (Calcein-AM-positive). Treatment with *O*^6^-benzylguanine reversed the protective effect of hAGT and overexpression of C145S-hAGT imparted minimal protection (Figure 1D). Similarly, live cell imaging with a viability dye Calcein-AM following PN treatment showed more number of Calcein-positive live cells only when hAGT was overexpressed (Figure 1E).

Other than cell death, DNA damage can trigger cellular stress responses such as senescence and polyploidy.[43] We were particularly interested in these two phenotypic changes because of their close associations with genomic instability seen in both tumor and aging cells.[44] Accordingly, we examined senescence marker SA-β-galactosidase (Figure 1F) and formation of polyploid cells (Figure 1G) following PN exposure of MDA-MB-231 cells in presence and absence of hAGT, C145S-hAGT and hAGT plus *O*^6^-benzylguanine. We found that hAGT indeed provided MDA-MB-231 cells appreciable protection from both (Figure 1F and 1G).

Together, these results indicated a novel role of hAGT and its active site Cys145 residue in providing protection from DNA cleavage and cytotoxicity caused by activated PBMC, induced macrophages and PN. It also confirmed a protective role from PN-associated growth arrest and genomic instability in target cells. To the best of our knowledge, this data presented the first evidence of a novel protective function of this well-studied ubiquitous repair protein hAGT.

### Overexpression of hAGT protects *E. coli* from the cytotoxic effect of PN

To determine if the cytoprotective function of hAGT is an inherent property of the protein, we examined its effect in a heterologous system. Accordingly, *E. coli* overexpressing hAGT or C145S-hAGT (Figure S6) were treated with PN because it is one of the major reactive intermediates formed during inflammation and also a major causative agent of neutrophil and macrophage-mediated cytotoxicity and DNA damage.[39] As expected, exposure of empty vector-transformed *E. coli* to PN (250 µM, 10 min on ice) followed by plating on a kanamycin LB agar plate resulted in a notable reduction in total number of colonies formed compared to the no treatment control (Figure 2A). In contrast, treatment of hAGT-overexpressing *E. coli* with PN had negligible cytotoxic effect as shown by formation of similar number of colonies to the no-treatment control (Figure 2A). In case of the C145S-hAGT-overexpressing *E. coli,* PN treatment caused less number of colonies compared to the no-treatment control (Figure 2A). Together, these data confirmed the protective effect of hAGT from the cytotoxic effect of PN and also indicated it as an inherent property of the protein. A point worth mentioning here is the small, yet reproducible protective effect observed with the C145S-mutated hAGT protein. This is not surprising, because there are multiple Cys residues in hAGT and as reported previously with GSH, those should be able to offer protection from the cytotoxic effect of PN, albeit much less efficiently compared to the active site reactive Cys145.

**Figure 2.**
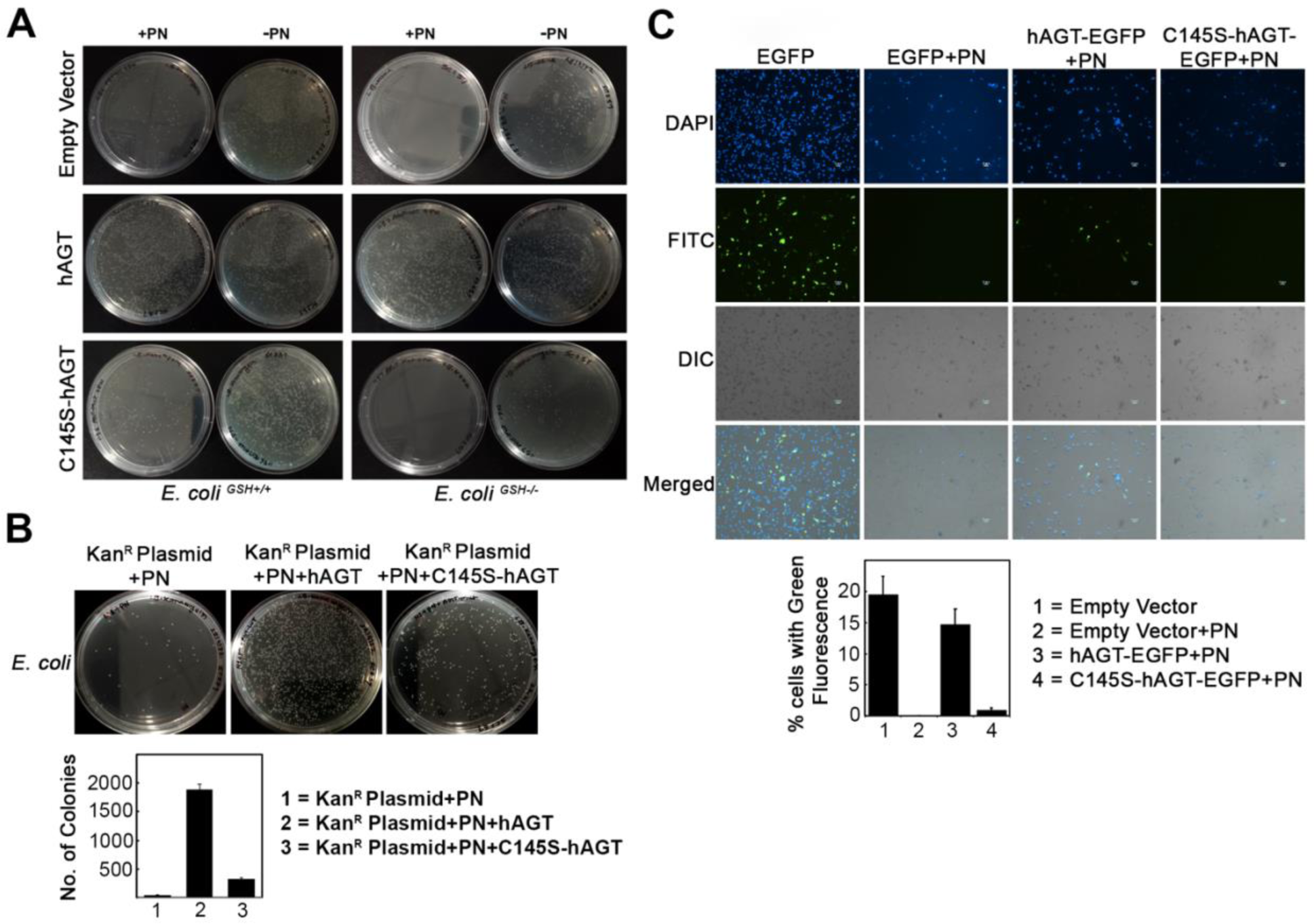
Protective effect of hAGT against PN-mediated damages is an intrinsic property of the protein. A) Kanamycin containing LB-agar plates showing colonies of *E. coli* that were transformed with either hAGT or C145S-hAGT expression construct containing the kanamycin-resistant selection marker and treated with PN after inducing the cells were IPTG. B) Kanamycin containing LB-agar plates showing colonies of *E. coli* that were transformed with kanamycin-resistance-conferring plasmid DNA after treating with PN in the absence or presence of hAGT or C145S-hAGT protein. C) MD-MBA-231 cells transfected with EGFP-expressing plasmid DNA that was pre-treated with PN in the presence of hAGT or C145S-hAGT protein. Images were acquired in Nikon Eclipse Ti microscope at 10X magnifications. Quantitation of fluorescence signals was done with ImageJ software. Each bar graph is an average of 3 readings with SE.

Because we have previously shown a similar protective role of GSH against the DNA damage and cytotoxic effects of PN,[21] we also repeated the above-mentioned experiments with homozygous GSH-negative strain of *E. coli* (*E. coli^GSH-/-^*). These results were similar to that obtained with the wild-type *E. coli* stain (Figure 2A).

### hAGT restores the PN-mediated loss of biological property of plasmid DNA

To determine the mechanism behind this novel protective function of hAGT, pEGFP-N1 plasmid DNA having the kanamycin-resistant gene was exposed to PN (250 µM) on ice for 10 min followed by incubation with buffer, hAGT or C145S-hAGT protein for 2 h at 37 °C.[45] Because of the recent discovery of DNA-stimulated nuclear proteases that digest DPCs into DNA-peptide crosslinks and our speculation of a possible DNA-hAGT crosslink formation,[34, 46] we digested the reaction mixture with trypsin before transforming it into *E. coli*. Transformed *E. coli* cells were plated on kanamycin-containing LB agar plates and incubated overnight at 37 °C. The functional integrity of the plasmid DNA was measured by its ability to confer kanamycin resistance to the transformed cells. We found greater number of colonies on the plates that were transformed with PN-damaged plasmid DNA treated with hAGT compared to the plates where the PN-damaged plasmid DNA was untreated or treated with C145S-hAGT (Figure 2B). Consistent with our earlier observations, the number of colonies on the C145S-hAGT-treated plates were more than the untreated control but significantly less than the hAGT-treated plate. We also performed parallel experiments with MDA-MB-231 mammalian cell line with the exception of trypsin digestion of the reaction mixture before transfection due to the presence of DNA-stimulated proteases in mammalian cells.[34] Here, the functional integrity of the pEGFP-N1 plasmid DNA was measured by its ability to produce green fluorescence through the expression of EGFP which renders the cells green when visualized with the FITC filter of an epifluorescence microscope. The results were similar to that obtained with the *E. coli* experiment (Figure 2C). Together these results indicated that the hAGT protein was able to restore the biological property of the plasmid DNA that was otherwise destroyed by PN. Such effect was primarily mediated through the active site C145 of hAGT. As already indicated, the minimal protection rendered by the C145S-hAGT protein could be due to other Cys residues within the protein. The possibility of direct neutralization of PN by hAGT was negated by adding hAGT 10 min after PN addition, which provided enough time for complete hydrolysis of PN.

### Effect of hAGT on PN-mediated DNA cleavage

Having demonstrated that hAGT can mitigate the effect of PN on plasmid DNA, we performed DNA cleavage assay to elucidate the mechanism of this novel function of hAGT. PN-mediated DNA damage results in the formation of various lesions including the major lesion 8-NO_2_-G.[23, 47, 48] 8-NO_2_-G being labile undergoes spontaneous depurination to form abasic sites which ultimately leads to DNA cleavage.[20] This process of converting PN-damaged sites in DNA into cleavage sites is facilitated by DMEDA workup. Typically, highly polymerized calf thymus DNA (ctDNA) produces a broad band in an agarose gel. Random DNA cleavage results in the generation of smaller fragments of various length that produces a smear in an agarose gel. The length of the DNA fragments is correlated with the extent of cleavage and can be used as a qualitative measure of DNA cleavage efficiency. Accordingly, ctDNA was treated with PN on ice for 10 min followed by addition of GSH, hAGT, C145S-hAGT or BSA (bovine serum albumin). Consistent with our published observation with GSH, PN caused significant DNA cleavage that was inhibited by the addition of GSH (5 mM) (Figure 3A). Addition of hAGT protein (5 μM) also provided similar inhibition of PN-mediated DNA cleavage. In contrast, both C145S-hAGT and BSA provided negligible protection from DNA cleavage. These results demonstrated that hAGT (5 μM) can either repair the PN-mediated DNA damage or similar to GSH (5 mM), prevents conversion of PN generated labile adducts into DNA strand breaks.

**Figure 3.**
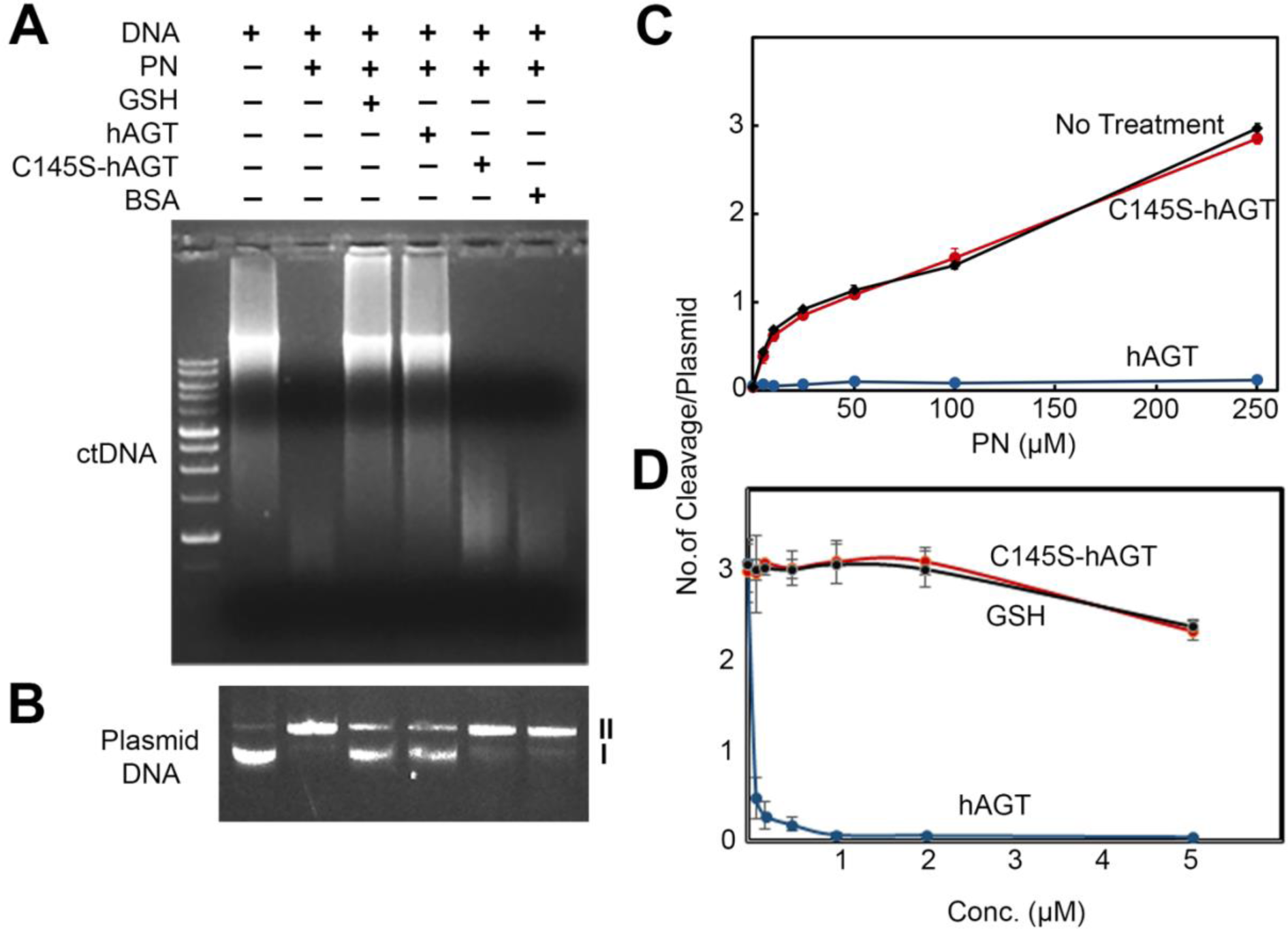
Effect of AGT on PN-mediated DNA cleavage in vitro. A) Agarose gel image of ctDNA after various treatments. B) Agarose gel image of pEGFP-N1 plasmid DNA after various treatments. C) pEGFP-N1 plasmid DNA cleavage by PN (0-250 μM) in the presence and absence of hAGT (5 μM) and C145S-mutated hAGT (5 μM). D) pEGFP-N1 plasmid DNA cleavage by PN (250 μM) in the presence and absence of varying concentration of hAGT (0-5 μM), C145S-hAGT (0-5 μM) or GSH (0-5 μM).

For quantitative analysis, we performed plasmid DNA cleavage assay as previously reported (Figure 3B).[49] We found that DNA cleavage by 0-250 μM of PN was completely inhibited by 5 μM of hAGT while similar concentration of C145S-hAGT had negligible effect, compared to the no treatment control (Figure 3C). Likewise, using varying concentration of the protein we found that 1 μM of hAGT was sufficient to completely inhibit DNA cleavage caused by 250 μM of PN while even up to 5 μM concentration of either C145S-hAGT or GSH has negligible effect (Figure 3D). Together these results demonstrated that the labile lesions formed in DNA due to PN treatment were either repaired or converted into non-labile lesions by hAGT and this process requires the presence of the active site Cys145.

### Effect of hAGT on PN-mediated cellular DNA cleavage

Next, we examined the ability of hAGT to mitigate PN-mediated genomic DNA cleavage in the complex environment of a cell. MDA-MB-231 cells, which has undetectable level of AGT (Figure S2), were treated with PN and the resultant genomic DNA cleavage of individual cells was determined using comet assay.[50] When MDA-MB-231 cells transfected with empty pEGFP-N1 vector were treated with PN, significant genomic DNA cleavage were observed as evident from the formation of comet tail (Figure 4A). No comet tail was observed upon treatment of hAGT-overexpressing MDA-MB-231 cells with PN while treatment of MDA-MB-231 cells-overexpressing C145S-hAGT protein produced comet tail similar to the empty vector control (Figure 4A). These data confirmed that hAGT can either repair PN mediated DNA damage or similar to GSH prevents conversion of PN-generated labile adducts into DNA strand breaks.

**Figure 4.**
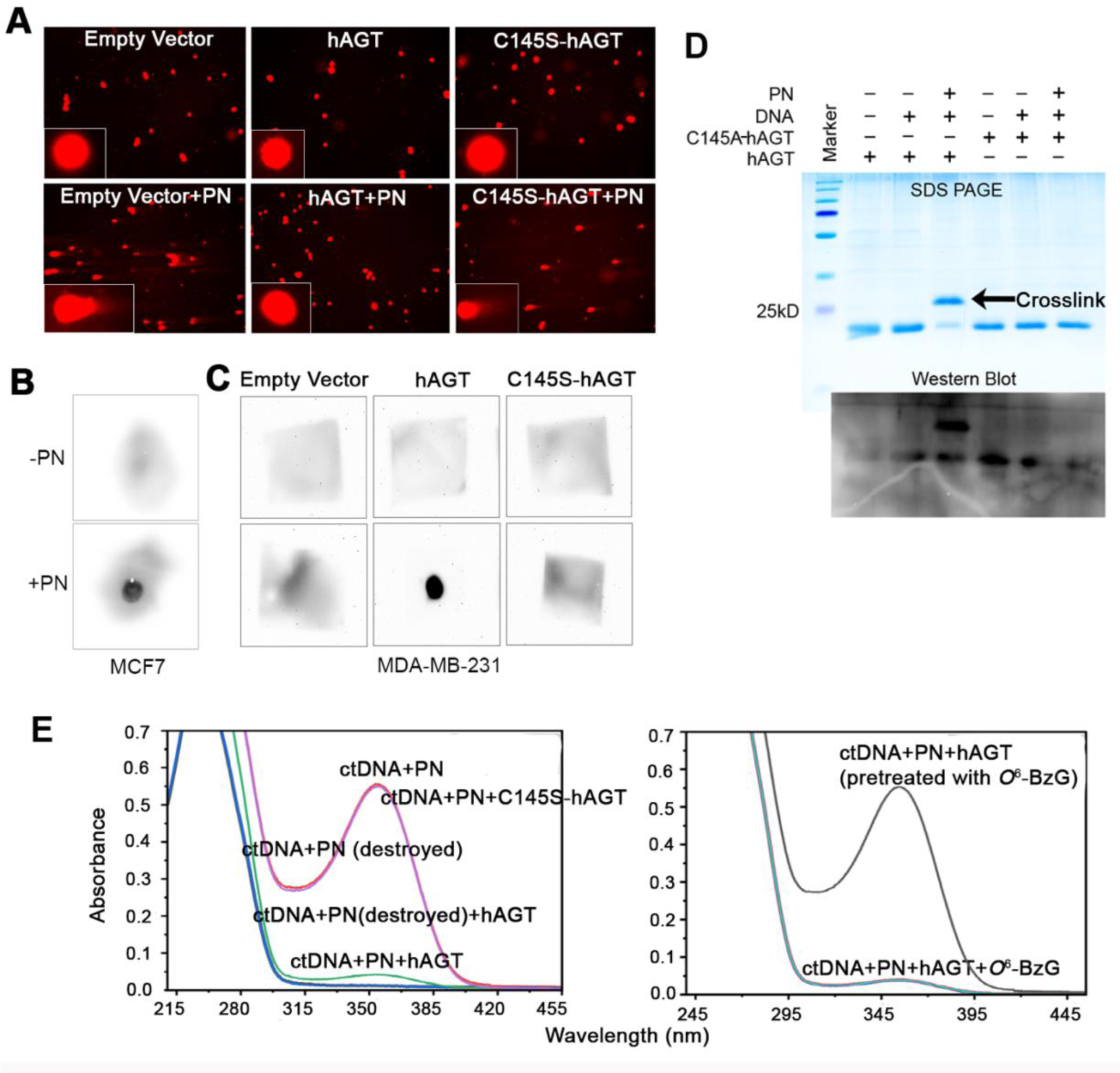
Inhibition of PN-mediated cellular DNA cleavage by hAGT and formation of DNA-AGT crosslink. A) Comet assay showing cellular DNA cleavage caused by PN and its inhibition by hAGT. Images were acquired at 10X (insert 20X) magnifications. B) Dot blot analysis showing covalently attached AGT to genomic DNA in MCF7 cells C) Dot blot analysis showing covalently attached AGT to genomic DNA in hAGT-overexpressing MDA-MB-231 cells. D) Gel shift assay showing covalent linkage between hAGT and a PN treated oligonucleotide. The presence of AGT-specific band in the Coomassie gel was further confirmed by western blot analysis. E) UV-Vis Spectra of ctDNA following various treatments.

### hAGT forms DNA-protein crosslinks (DPC) when cells are exposed to PN

To determine if this novel protective function of hAGT from PN-mediated DNA cleavage is either due to repair or conversion of PN-generated labile adducts into stable adducts, we performed a dot blot assay. MCF7 cells, due to its high level of endogenous hAGT expression (Figure S2), were treated with PN (2.5 µM) for 4 min in ice cold PBS, washed and incubated for 3 h. Because hAGT has a binding affinity for DNA, non-covalently-bound proteins were stripped from DNA by using a published protocol by adding 200 mM KCl and shearing the genomic DNA by repeated pipetting.[51] This process was repeated three times to ensure complete removal of any non-covalently-bound proteins. The DNA pellet obtained from the reaction mixture was finally subjected to dot blot analysis using anti-hAGT antibody. While negligible signal was observed on the blot for the untreated cells, treatment with PN produced a strong signal (Figure 4B). Similarly, when the experiment was repeated with MDA-MB-231 cells, a strong signal was observed only for the PN-treated hAGT-overexpressing cells (Figure 4C). Together, these data indicated the formation of DNA-hAGT crosslink in PN-treated cells. Although the DPC formation does not provide any direct evidence regarding the mechanism behind the novel protective function of hAGT from PN-mediated DNA cleavage, it is consistent with our previously proposed mechanism with GSH. Conversion of the labile 8-nitroguanine DNA lesion into a stable DPC prevents DNA cleavage and the toxicity resulting from it.

### hAGT forms DNA-protein crosslinks with PN-damaged DNA

To confirm the ability of hAGT to form DPC with PN-damaged DNA, we performed a gel-shift assay (Figure 4D).[52] DNA gel-shift assays are widely accepted for the *in vitro* detection of DPC. To exclude non-covalent DNA binding of proteins, the gel-shift assay was performed under denaturing conditions. A 120 bp DNA fragment was treated with PN (2 µM) on ice for 10 min before addition of 600 ng purified recombinant hAGT or C145S-hAGT protein. After incubation of the reaction mixture at 37 °C for 2 h it was resolved in 15% PAGE under denaturing conditions. In a separate experiment western blot analysis was performed to confirm the identity of the DNA-hAGT crosslink band. As expected, purified hAGT yielded a band at∼23 kD both in the absence and presence of DNA (Figure 4C). However when the DNA was pre-treated with PN before addition of hAGT, majority of the ∼23 kD band shifted to ∼26 kD. In case of the C145S-hAGT protein, no such shift was observed. The identity of the shifted band was confirmed by western blot analysis. These results clearly showed that hAGT forms a stable covalent linkage with PN-damaged DNA through its active site Cys145.

To further confirm the formation of the DNA-hAGT crosslink in PN-treated DNA, we performed a UV-Vis experiment (Figure 4E). It has been shown that 8-NO_2_-G-containing DNA has a distinct peak at 354 nm apart from the normal 260 nm peak for DNA.[53] If reaction of hAGT with 8-NO_2_-G in DNA follows a similar mechanism as proposed for GSH, then formation of DNA-hAGT crosslink should result in removal of the nitro group from 8-NO_2_-G in DNA and consequently the 354 nm peak should disappear. Indeed when ctDNA was treated with 5 mM PN, a peak at 354 nM was observed which was not present in ctDNA incubated with PN pre-treated with neutral buffer (this process destroys PN). The 354 nM peak almost disappeared upon addition of hAGT (5 µM). Addition of 1-5 µM of *O*^6^-benzylguanine (which forms a covalent adduct with hAGT) to the DNA+hAGT reaction mixture did not produce any change indicating removal of 8-NO_2_-G from the DNA. When hAGT (5 µM) was pre-treated with *O*^6^-benzylguanine (5 µM) and then added to the PN treated ctDNA, the 354 nm peak remained unaltered, suggesting that inactivated hAGT cannot remove 8-NO_2_-G from the DNA. Similarly, when the C145S-hAGT was added there was no effect on the 354 nm peak, indicating that the Cys145 thiol is essential for the removal of 8-NO_2_-G from the DNA. Together, these results indicated that hAGT forms a covalent adduct with 8-NO_2_-G in DNA by a mechanism that involves loss of the nitro group and formation of a AGT-DNA crosslink, possibly by nucleophilic substitution of the nitro group by Cys145 thiol of hAGT.

### Activated macrophages cause induction of hAGT in mammalian cells

Because the DNA-hAGT crosslink formation requires stoichiometric amounts of hAGT, we examined if the oxidative and nitrosative stress associated with inflammation has the ability to induce hAGT expression in participating immune cells as well as host cells. Accordingly, we co-cultured induced macrophages with MDA-MB-231 cells (express undetectable levels of endogenous hAGT, Figure S3) in transwell chambers and performed qRT-PCR for AGT transcript and western blot for AGT protein at various time points. We found a time-dependent (24 and 36 h) increase in both transcript and protein levels of hAGT in both macrophages and MDA-MB-231 cells (Figure 5A-B). We also looked into the effect of the anti-inflammatory agent quercetin on macrophage-mediated induction of hAGT. As expected, we found that addition of quercetin significantly reduced hAGT induction, as depicted by western blot analysis (Figure 5C). This indicates that inflammation itself is a trigger for hAGT expression, possibly to render its protective effect on DNA. The mechanistic detail behind this interesting, yet preliminary observation is unclear and out of scope of this manuscript. Epigenetic regulation of hAGT expression is a widely-studied topic, particularly in the context of transcription factors AP1, SP1, NF-κΒ, C/EBP as well as DNA and histone methylation. Majority of these factors are redox-modifiable.[54, 55] Although hAGT is significantly more efficient than the known protective agent GSH in protecting the cell from the cytotoxic effect of PN during inflammation, one caveat is the high cellular concentration of GSH (5-10 mM) and the lower base-line concentration of hAGT. The finding that the hAGT expression is induced by inflammation clearly indicates that the difference in cellular concentration between GSH and hAGT may not be an issue in inflamed tissue.

**Figure 5.**
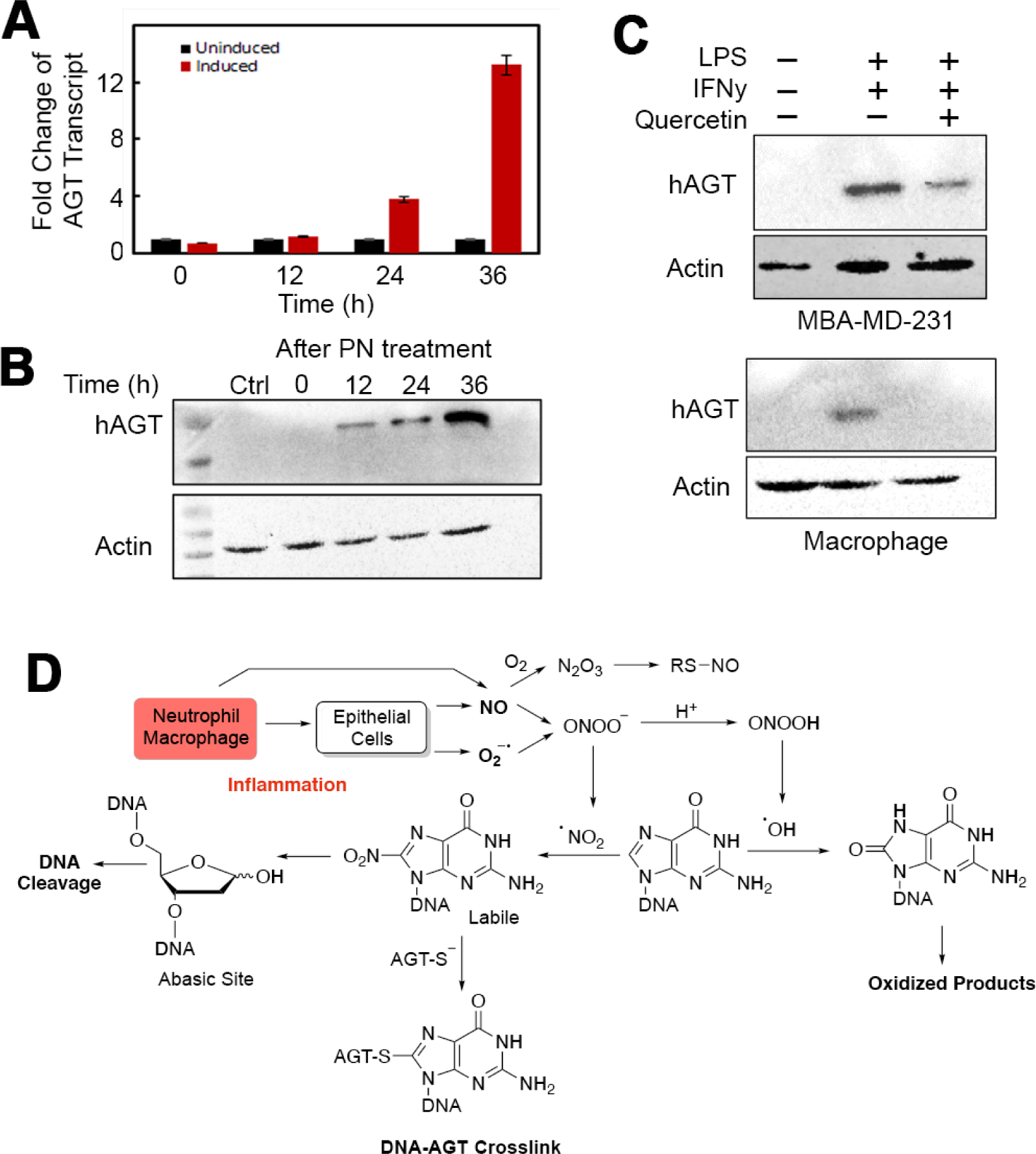
Induction of hAGT expression by inflammation mediated reactive species. A) Induction of hAGT transcript in MDA-MB-231 cells in the presence of induced macrophages. Using qRT-PCR fold change in hAGT transcript were detected in MDA-MB-231 cells that were cocultured with induced macrophages in transwell chambers. Each bar graph is an average of 3 readings with SE. B) Western blot analysis showing induction of hAGT protein in MDA-MB-231 cells after various time interval following treatment with PN for 4 min. C) Effect of the anti-inflammatory agent quercetin on macrophage-mediated induction of hAGT in both the target MDA-MB-231 cells and induced macrophages. D) Proposed pathway for the formation of 8-nitroguanine and 8-oxoguanine in DNA during inflammation.

### Proposed Mechanism

Based on our previous work on the GSH-DNA adduct,[21, 47, 56] we proposed a mechanism for the formation of the DNA-hAGT crosslink as shown in Figure 5D. The proposed mechanism involves nucleophilic attack by the Cys145 thiolate of hAGT (pK_a_ ∼5) on the C8 atom of 8-NO_2_-G followed by elimination of the NO_2_ group, leading to the formation of a stable DNA-hAGT crosslink. Here, hAGT being a protein (and not an enzyme) can perform the DPC-forming chemistry only once with each 8-NO_2_-G lesion. Hence, stochiometric amounts of hAGT is needed during inflammation to compete with the 8-NO_2_-G lesions formed in DNA. Strikingly, hAGT was found to be induced by both PN and activated macrophages. The ability of the cell to stimulate the *de novo* synthesis of hAGT during inflammation warrants that this genome-maintaining property is an important and essential function of this protein.

The hAGT-DNA crosslink formation as a genome-maintenance strategy seems counterintuitive and detrimental for the cell, and a heavy price to pay for the protection it provides from the toxicity inflicted by PN-mediated DNA damage. This is because DPCs are bulky lesions that present a challenge for the cellular repair systems and often thought to cause replication block. However, the presence of DNA-stimulated proteases that can digest bulky DPCs[33, 34], translesion polymerases that can bypass DNA-peptide crosslinks,[33] and ability of the cell to repair DNA-peptide crosslinks, probably through NER and HR,[57] suggest otherwise. In fact, the hAGT-DNA crosslink provides a way by which the PN-generated 8-NO_2_-G lesions are shielded from being spontaneously converted into abasic sites and strands breaks by the cell, till repair sets in. Although the mechanism of maintenance of genome integrity by forming DPC may seem unusual, the HMCES protein is known to have a similar function with respect to abasic sites.[58]

### Conclusions

In conclusion, we have discovered a novel function of the repair protein hAGT involving protection from the cytotoxicity and stress-response caused by PN-mediated DNA cleavage. Using a custom designed “transwell assay” with activated PBMC or induced macrophages and various cell culture assays with hAGT or C145S-hAGT-overexpressing mammalian cells, we have demonstrated that hAGT can protect cells from the genomic instability and cytotoxicity caused by PN-mediated DNA damage during inflammation. We have further shown that this protection is due to formation of a stable hAGT-DNA crosslink. With the help of C145S-hAGT and the hAGT inhibitor *O*^6^-benzylguanine, we also have provided evidence that the active site Cys145 thiolate of hAGT is involved in crosslink formation. Finally, we have demonstrated that induced macrophages can enhance the expression of hAGT. hAGT being a protein (and not an enzyme) with DNA binding affinity, presence of active site thiolate and inducible by PN and immune cells, has all the necessary properties to fulfil the previously unrecognized role of genome maintenance and protection of the cell from inflammation-associated toxicity. It is our inference that hAGT is a dual function protein that along with DNA repair, maintains genome integrity by forming hAGT-DNA crosslink.

## Materials and Methods

### Safety Statement

Peroxynitrite (PN) is extremely reactive at neutral pH, cytotoxic and a potential mutagen. Hence, it should be handled with extreme care and destroyed by adding neutral aqueous buffer before discarding.

### Reagents and Cell Lines

Unless otherwise mentioned, all reagents were of the highest purity available and obtained from either Sigma-Aldrich (now Merck India, Bangalore, India) or HiMedia Laboratories (Mumbai, India). N,N′-dimethylethylenediamine (DMEDA) was procured from Tokyo Chemical Industry (Tokyo, Japan); acetonitrile from JT Baker (Center Valley, PA, USA) and glutathione from Sisco Research Laboratories Pvt. Ltd. (Mumbai, India). Fluorescent dyes were purchased from Thermo Fisher Scientific (MA, USA)MDA-MB-231, MCF-7 and THP-1 cell lines were procured from National Center for Cell Sciences (Pune, India). Healthy human PBMCs were obtained from HiMedia Laboratories. The *E. coli^GSH-/-^* was a gift from Dr. A. K. Bachhawat (IISER-Mohali, Punjab, India).

### Synthesis of PN

PN was prepared using a published protocol and before each reaction its concentration was determined.[59] Briefly, following three solutions were freshly prepared in distilled water: 0.7 M HCl + 0.6 M H_2_O_2_, 0.6 M sodium nitrite, 3 M sodium hydroxide and taken into individual syringes. Contents of the individual syringes were mixed and immediately quenched by collecting in a beaker containing 3 M NaOH on ice. Concentration of PN was determined by taking its absorbance at 302 nm wavelength (extinction coefficient 1670 M^-1^ cm^-1^).

### Cloning

The full-length cDNA encoding hAGT (wild type/WT) and C145S-hAGT (mutated) were originally obtained from Prof. F. P. Guengerich (Vanderbilt University Medical Center, Nashville, TN, USA). These were cloned into pET30b (for recombinant protein expression) and pEGFP-N1 (for mammalian expression) vectors using standard protocol. The clones were confirmed by restriction digestion and Sanger sequencing.

### Recombinant WT and C145S-hAGT expression and purification

WT and C145S-hAGT cDNA in pET30b vector were transformed into *E. coli* (Rosetta) cells (a gift from Dr. Rajan Vyas, Shiv Nadar Institution of Eminence, Greater Noida, India) following standard protocols. A single transformed colony was picked up and inoculated in LB medium containing 50 µg/ml kanamycin at 37°C. When the optical density at 600 nm of the culture reached 0.6, cells were induced with 1 mM IPTG and grown for another 6 h at 30 °C. Cells were pelleted by centrifugation at 6000 g for 15 min and stored at -70°C for further use. For protein purification, the cell pellet was resuspended in lysis buffer (20 mM Tris-HCl pH 7.5, 150 mM NaCl, 5 mM imidazole) and sonicated for 10 min with a pulse cycle of 5 sec on, 20 sec off, 40% amplitude on ice followed by centrifugation at 22,500 g for 30 min. The supernatant was passed through a 0.45 µM syringe filter before transferring into a tube containing HIS-Select Nickel Affinity Gel (Merck India) and kept on a mixer at 4°C for 3 h. The mixture was centrifuged and the pellet was washed 3 times with 3 bed volumes of resuspension buffer (20 mM Tris-HCl pH 7.5, 150 mM NaCl, 20 mM imidazole). The His-tagged hAGT and C145S-hAGT proteins were eluted by 50 mM Tris-HCl pH 7.0, 150 mM NaCl including 300 mM imidazole. The purity of the proteins was examined by SDS-PAGE and the concentration was measured with Bradford reagent (Sigma-Aldrich).

### PBMC

Human peripheral blood mononuclear cells (PBMC) from single donor were obtained from HiMedia laboratories and activated with 100 ng/ml of LPS for 15 min as per the published protocol.[60]

### THP-1 monocyte/macrophage

THP-1 monocytes (NCCS, Pune, India) were differentiated into macrophages with 150 nM phorbol myristate acetate/PMA treatment for 24 h, washed with PBS and kept in RPMI, 10% FBS and 1% Pen-strep. Within five days cells were differentiated into macrophages and became adherent. These were further treated with 100 ng/ml LPS and 35 ng/ml of IFN γ for M1 polarization for 48 h. The differentiation status was verified with macrophage differentiation markers.[61]

### Co-culture Experiment

MDA-MB-231 cells were transfected with empty pEGFP-N1 or with pEGFP-N1-hAGT or pEGFP-N1-C145S-hAGT vector (3 µg each). Transfected cells were labelled with Cell-tracker™ Blue CMAC (25 μM; Thermo Fisher Scientific). PMA-treated THP-1 macrophages induced with LPS + IFNγ were labelled with Cell-tracker™ Orange CMRA (2 μM; Thermo Fisher Scientific). LPS-treated PBMCs were labelled with Cell Tracker™ Green CMFDA (20 μM; Thermo Fisher Scientific). Labelling was done in serum free media for 2 h. Labelled cells were trypsinized and co-cultured in 24 well plate (1.25 x 10^4^ cells of immune cells and 2.5 x 10^4^ cells of MDA-MB-231). Images were taken at 0, 12 and 36 h in appropriate fluorescent channels (Leica DFC450, Leica Microsystems, Wetzlar, Germany). To examine reactive oxidant production, induced PBMCs (in co-culture with MDA-MB-231) were exposed to MitoSOX Red™ mitochondrial superoxide indicator (Thermo Fisher Scientific) and images were captured at 0, 1 and 6 h.

### Transwell Assay

Transwell assays were performed using 8.0 μM pore, 6.5 mm polycarbonate transwell filters (HiMedia Laboratories). 1.25 x 10^4^ THP-1 macrophages polarized to M1 were seeded into the transwell filters fitted into 24-well plate. In the bottom well of the 24-well plate, 2.5 x 10^4^ MDA-MB 231 cells transfected with pEGFP-N1, hAGT, C145S-AGT were plated. Cells were maintained in serum free DMEM for 6 hours for comet assay and qRT-PCR.

### Measurement of Cell Viability and Cytotoxicity with Calcein-AM, PI and Annexin V live cell microscopy

First, hAGT and C145S-hAGT plasmids were transfected into MDA-MB 231 cells by using PEI as mentioned elsewhere. After 6 h, hAGT-transfected cells were exposed to 100 μM *O*^6^-benzylguanine. At 24 h post-transfection, all the transfected groups as well the non-transfected control were treated with 2.5 μM PN along with *O*^6^-benzylguanine in appropriate combinations in ice-cold PBS for 4 min. After the treatment, cells were washed with 1X PBS twice and kept in fresh complete media for another 48 h. Images were taken in Leica DFC450C microscope by staining cells with viability marker Calcein-AM, death marker PI and apoptosis marker Annexin V (all from Thermo Fisher Scientific).

### SA-β-galactosidase Assay

hAGT and C145S-hAGT plasmids were transfected in MDA-MB 231 cells as mentioned elsewhere. After 6 hours, hAGT-transfected MDA-MB-231 cells were pretreated with 100 μM *O*^6^-benzylguanine. After 24 hours of transfection, all transfected groups as well the non-transfected control were exposed to 2.5 μM PN along with *O*^6^-benzylguanine in one of the hAGT-transfected groups which was pretreated earlier for 4 min in ice-cold PBS. After the treatment, cells were washed with 1X PBS twice and fresh complete media was added. After 48 hours of PN treatment, β−gal staining solution was added to each group according to previously published protocol.[62]

### Assessing Polyploidy and micronuclei

hAGT and C145S-AGT plasmids were transfected in MDA-MB 231 cells as mentioned elsewhere. After 6 hours of transfection, hAGT-transfected MDA-MB-231 cells were pretreated with *O*^6^-benzylguanine. After 24 hours of transfection, all the transfected groups as well the non-transfected control were exposed to 2.5 μM PN along with 100 μM *O*^6^-benzylguanine in one of the hAGT-transfected sets which was pretreated earlier. PN treatment was done in ice-cold PBS for 4 min. After PN exposure, cells were washed with 1X PBS twice and added to fresh, complete media. After 48 h PN treatment, 500 nM Hoechst stain (Thermo Fisher Scientific) was added to each set of experiments and images were captured with Leica DFC450.

### Effect of hAGT overexpression on the cytotoxicity of PN in *E. coli*

*E. coli^GSH+/+^* and *E. coli^GSH-/-^* competent cells were transformed with empty pET30b, hAGT pET30b or C145S-hAGT pET30b plasmid expression constructs and grown on kanamycin-containing LB agar plate overnight at 37°C. Single colony was picked up and cultured in LB media containing 50 µg/ml kanamycin. When the OD at 600 nm reached 0.6, culture was induced by adding 1 mM IPTG. Following 5 h induction at 30°C, cells were pelleted by centrifugation at 3000 rpm, washed and resuspended in PBS. Resuspended cells were treated with 250 µM PN on ice for 10 min before plating on kanamycin containing LB agar plate. The plates were incubated overnight at 37°C.

### Effect of hAGT on PN-damaged plasmid DNA

pEGFP-N1 plasmid DNA (50 µg) in 100 mM potassium phosphate (KPhos, pH 7.5) buffer was treated with PN (250 µM) on ice for 10 min before addition of hAGT or C145S-hAGT (5 µM) recombinant protein and incubated at 37°C for 2 h. A control reaction with no hAGT added was also kept. After incubation, the product was digested overnight with trypsin at 37°C followed by precipitation of the DNA using 70% ethanol and 0.3 M sodium acetate. The precipitated plasmid DNA was pelleted by centrifugation at 10000 g, washed with cold 70% ethanol, air-dried and finally transformed into competent *E. coli* cells following standard protocol. The transformed cells were plated on LB agar plates containing kanamycin (50 µg/mL), incubated overnight at 37°C and colonies were manually counted. The reactions were carried out in triplicate.

To determine the effect in mammalian cells, the DNA, after treatment with PN and hAGT, was precipitated using 70% cold ethanol and 0.3 M sodium acetate. The precipitated plasmid DNA was pelleted by centrifugation at 10000g, washed with cold 70% ethanol, and air-dried. MDA-MB-231 cells were transfected with precipitated plasmid (3 µg) using polyethylenimine (PEI), at 2:1 PEI:DNA ratio. Imaging was done to determine the expression of EGFP after 48 h of transfection using a Nikon Eclipse Ti microscope.

### DNA Cleavage Assay

In a typical DNA cleavage reaction, pEGFP-N1 plasmid DNA (1 µg) in KPhos buffer (100 mM, pH 7.4) was treated with PN (0−250 µM) on ice for 10 min followed by addition of 5 µM of hAGT or C145S-hAGT recombinant protein and incubated at 37 °C for 2 h. For assays involving the effect of hAGT on DNA cleavage by PN, pEGFP-N1 plasmid DNA in KPhos buffer (100 mM, pH 7.4) was treated with PN (250 µM) on ice for 10 min followed by the addition of 0-5 μM hAGT or C145S-hAGT protein. Following incubations, reactions were subjected to N,N′-dimethylethylenediamine (DMEDA, 100 mM) workup for 2 h at 37°C, quenched by the addition of 5 µL of glycerol loading buffer, and then electrophoresed for 60 min at 100 V in 1% agarose gel (w/v) containing 0.5 µg/mL ethidium bromide. DNA was visualized and quantified using a gel documentation (Bio-Lad Laboratories, Hercules, CA, USA) system. Strand breaks per plasmid DNA molecule (S) were calculated using the equation S = −ln f I, where f I is the fraction of plasmid present as form I.

For assays involving calf thymus (ct) DNA, 50 µg of ctDNA was treated with 2 mM PN at 25°C for 10 min followed by addition of hAGT (5 µM), C145S-hAGT (5 µM), BSA (5 µM) or GSH (5 mM). Labile sites were converted into cleavage sites using DMEDA workup and analysed by agarose gel electrophoresis. The extent of DNA cleavage was visualized by the length of DNA smear in an agarose gel.

### Comet Assay

Comet assay was performed following the published protocol of Dhawan et al.[63] Briefly, 5 x 10^5^ MDA-MB-231 cells were seeded in 35 mm plate and transfected with empty pEGFP-N1 plasmid or with pEGFP-N1 plasmid having either hAGT or C145S-hAGT cDNA (3 µg) using polyethyleneimine at 2:1 PEI: DNA ratio. After 48 h of transfection, cells were transferred into ice cold PBS and treated with PN (5 µM) for 4 min. After PN treatment cells were washed with PBS and kept for 1 h in complete media. 1.5 × 10^4^ PN-treated transfected cells were added into 0.5% low melting agarose (LMPA) and layered on a 1% agarose-coated glass slide. Embedded cells were lysed in lysis buffer containing 2.5 M NaCl, 100 mM EDTA, 10 mM Tris-HCL, pH 10 and TritonX-100 followed by DNA unwinding in alkaline buffer (300 mM NaOH and 1 mM EDTA, pH >13) for 30 min. Slides were electrophoresed in the same alkaline buffer at 21 V for 40 min. Cells were neutralized with 400 mM Tris-HCl, pH 7.5 and stained with ethidium bromide (2 µg/mL) and visualized using a Leica DFC450C microscope.

### Detection of DNA-hAGT crosslink formation in cells by dot blot analysis

1.5 x 10^6^ MCF7 or MDA-MB-231 cells (transfected with empty pEGFP-N1 plasmid or with pEGFP-N1 plasmid having either hAGT or C145S-hAGT cDNA), were treated with 2.5 µM PN for 4 min in ice-cold 1X PBS and then washed twice with PBS. Cells were then incubated for 3 h in complete media, lysed with 0.5 ml of 2% SDS containing 1 mM PMSF and 20 mM Tris-HCl (pH 7.5) and stored at -70°C. For dot blot analysis, the mixture was thawed, vortexed at high speed for 10 sec and incubated at 65°C for 10 min. 0.5 ml of 200 mM KCl in 20 mM Tris-HCl was added and the mixture was passed through 1 ml pipette tip 5-8 times to shear the DNA. The reaction mixture was kept on ice for 5 min and centrifuged at 3000 g for 5 min at 4 °C. The pellet was resuspended in 1 ml of 100 mM KCl, 20 mM Tris-HCl and again sheared by pipetting. The last two steps were repeated twice. Finally, the pellet was resuspended in 1 ml H_2_O and proceeded for dot blot analysis with the anti-AGT antibody (Cell Signalling Technologies, Danvers, MA, USA).

### Gel Shift Assay

A 120-bp PCR-amplified DNA fragment (5 μM) was incubated with 2 µM PN in 100 mM KPhos buffer (pH 7.5) on ice for 10 min, followed by the addition of 600 ng of hAGT or C145S-hAGT protein. The reaction mixture was incubated at 37°C for another 2 h and subjected to 15% SDS-PAGE analysis. Proteins were visualized by either Coomassie staining or western blot analysis.

### Detection of DNA-hAGT crosslink formation in ctDNA by UV-Vis analysis

Calf thymus (ct) DNA (50 μg) in potassium phosphate buffer (100 mM, pH 7.4) was treated with PN (5 mM) for 10 min on ice followed by UV-Vis analysis. A peak at 355 nm was observed corresponding to the formation of 8-NO_2_-G. To the PN-treated ctDNA, 5 μM hAGT, C145S-hAGT or 5 μM *O*^6^-benzylguanine pre-treated hAGT was added and incubated for 30 min at 37°C before collecting the UV-Vis spectra. In a separate experiment, PN-treated ctDNA exposed to hAGT was treated with various concentration of *O*^6^-benzylguanine (1-5 μM) and incubated for 5 min at 37°C before collecting the UV-Vis spectra. Absorbance was recorded in a Shimadzu UV-2600i UV-VIS Spectrophotometer.

### QRT-PCR Analysis

Total RNA was extracted from cells using Trizol (Thermo Fisher Scientific), treated with TURBO DNA-*free*™ Kit (Thermo Fisher Scientific), quantified using NanoDrop (Thermo Fisher Scientific). QRT-PCR reactions were performed by using the GoTaq®1-Step RT-qPCR System (Promega Scientific, Madison, WI, USA) with Bio-Rad CFX Connect (Bio-Rad Laboratories) instrument. Actin was used as house-keeping control. Relative mRNA expression levels were calculated by the standard deltaCt method.[62]

### Quercetin Treatment followed by western blot analysis of LPS+IFNγ−induced M1 macrophages

THP-1 monocytes were treated with 150 nm PMA for 24 hour to differentiate into macrophages. After 5 days of differentiation, macrophages were co-treated with 1 μg LPS and 35 ng of IFNγ along with 39 μg/ml quercetin. Induced macrophages were kept inside a transwell chamber and MDA-MB-231 cells kept in the outside chamber. After 36 h, protein was collected from both macrophages and MDA-MB-231 cells for western blot analysis using anti-AGT antibody.

### Western blot and qRT-PCR of PN-treated MDA-MB 231 cells

MDA-MB-231 cells were treated with 2.5 μM PN in ice-cold PBS for 4 min. Cells were subsequently washed twice with 1X PBS before adding fresh complete media. After incubating the PN-treated MDA-MB 231 cells for 0, 12, 24 and 36 h, protein and RNA were isolated for western blot and qRT-PCR PCR for hAGT.

### Western blot

Isolated proteins were denatured using 5X denaturation buffer at 95 °C for 10 min and then resolved on 12% SDS-PAGE. Resolved proteins were transferred onto a nitrocellulose membrane and blocked by incubating in 5% skimmed milk. Blots were further incubated with primary antibodies overnight at 4℃, followed by washing three times (10 min each) with 1X Tris buffer saline having 0.1% Tween-20 (TBST) before incubation with HRP-conjugated secondary antibody. Unbound antibodies were removed by washing with 1X TBST buffer (3×10min) and chemiluminescence signal was recorded using Pierce ECL Western Blot Substrate (Thermo Fisher Scientific) with Bio-Rad ChemiDoc (Bio-Rad laboratories). Blots were stripped and incubated with anti β-actin antibody as internal control.

## SUPPORTING INFORMATION

Supporting Information is available. The file contains material and methods, fluorescence microscopic images and protein gel images.

## AUTHOR CONTRIBUTION

Conceptualization, G.C.; Methodology, G.C., A.C.; Investigation, S.C., G.M., A.C., and G.C.; Writing-original draft, G.C. and AC; Writing-review and editing, G.C., A.C.; Funding acquisition, A.C. and GC; Supervision, A.C. and G.C.

## ACKNOWLEDGEMENT

This work was supported in part by Shiv Nadar Institution of Eminence [to A.C.]. The authors also thank Prof. F. P. Guengerich for providing with the original hAGT construct and Prof. Bacchawat for providing the *E. coli^GSH-/-^ cells*.

---

**Table of Content Figure.**
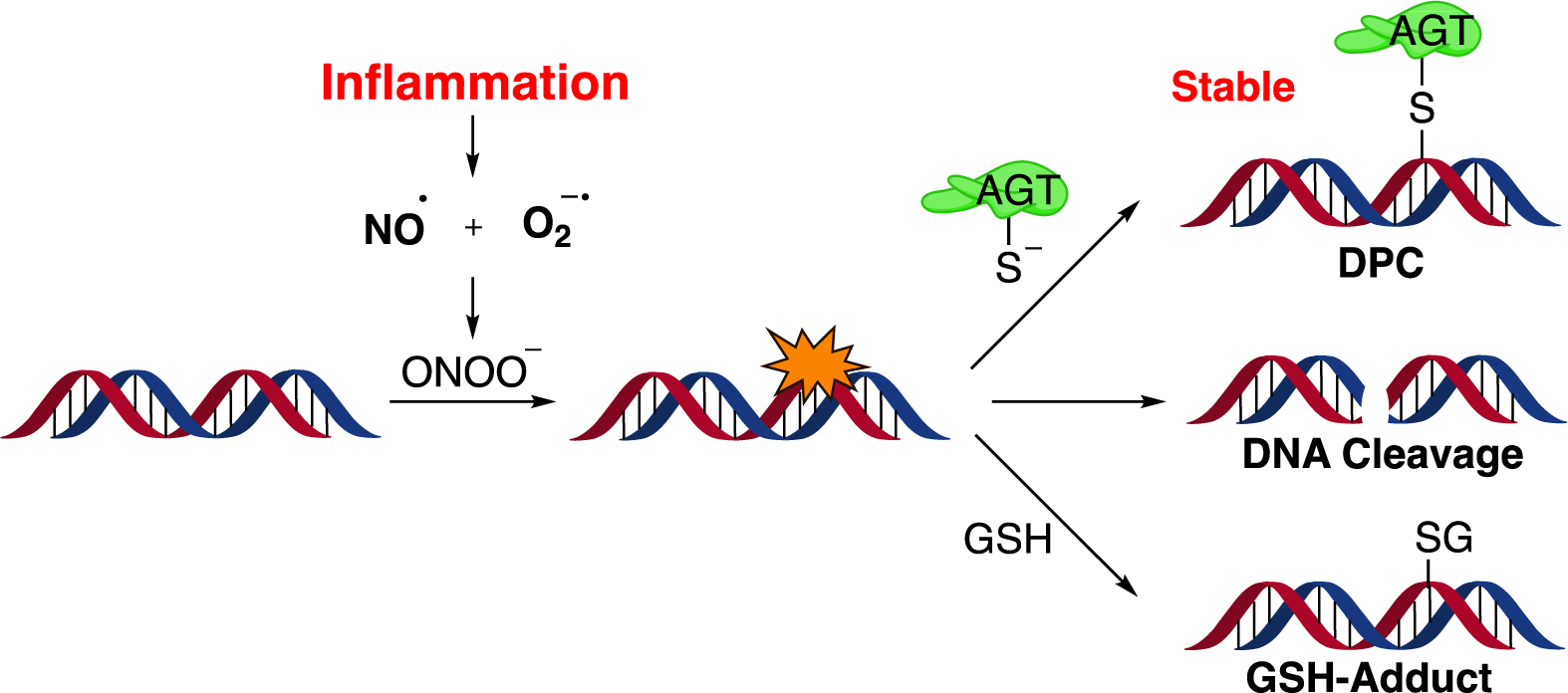

Peroxynitrite (PN), generated during inflammation cause DNA damage including 8-nitroguanine formation. Here, we report the discovery of a protective function of the repair protein *O*^6^-alkylguanine-DNA alkyltransferase (hAGT/MGMT). AGT was found to maintain genome integrity by forming DNA-protein crosslink.

