## Supplemental Data for "*O*^6^-Alkylguanine-DNA Alkyltransferase Maintains Genomic Integrity During Peroxynitrite-Mediated DNA Damage by Forming DNA-Protein Crosslinks"

**Contents:**

**Figure S1:** Detection of mitochondrial ROS formation in the co-culture of MDA-MB-231 cells and activated PBMC for 0, 1 and 6 h.

**Figure S2:** Effect of hAGT on activated PBMC-mediated clearance of target epithelial cells.

**Figure S3:** Detection of hAGT or C145S hAGT in the cell lysate of MDA-MB-231 cells transfected with hAGT pEGFP-N-1 or C145S-hAGT pEGFPN-1.

**Figure S4**: Differentiation of THP1 monocyte to macrophage.

**Figure S5:** Effect of hAGT on induced macrophage-mediated clearance of target epithelial cells.

**Figure S6:** Coomassie stain images of recombinant and purified hAGT and C145S hAGT.

**Figure S1.** MDA-MB-231 cells (labelled with cell tracker blue) were co-cultured with activated PBMC for 0, 1 and 6 h and analysed for reactive species generation using MitoSox Red reagent by formation of red fluorescence.

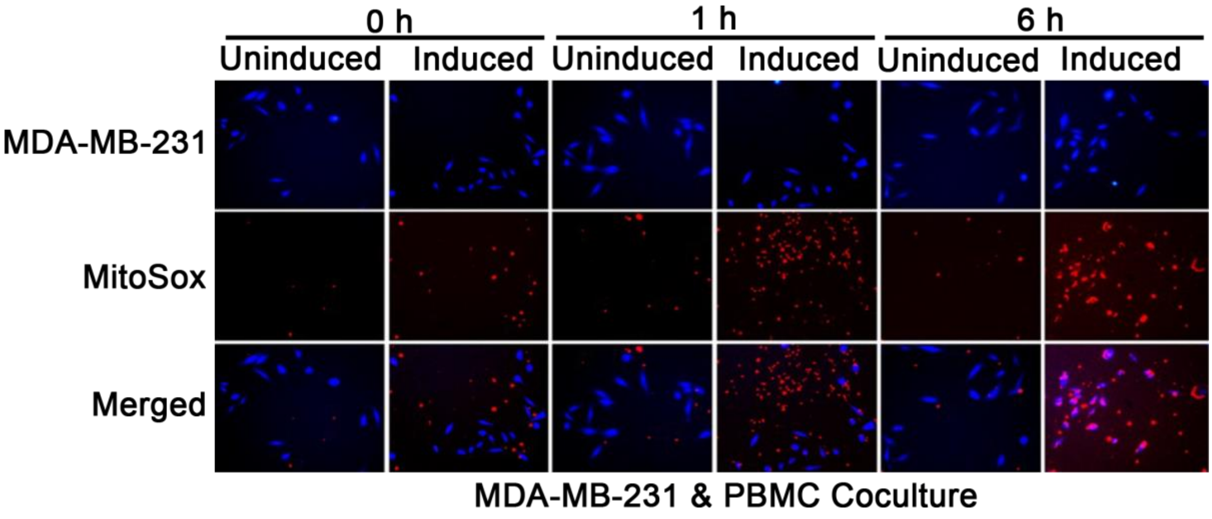

**Figure S2.** Effect of hAGT on activated PBMC-mediated clearance of target epithelial cells. Live cell images of co-cultured MDA-MB-231 and activated PBMCs at A) 0 h and B) 36 h. MDA-MB-231 cells were labelled with cell tracker blue and PBMCs were labelled with cell tracker green. MDA-MB-231 cells were transfected with empty pEGFP-N1, hAGT pEGFP-N1 or mutated-C145S hAGT pEGFP-N1 vectors for overexpression of the corresponding proteins and before co-culturing with activated PBMCs. In one set of experiments, hAGT pEGFP-N1-expressing MDA-MB-231 cells were also treated with the known inhibitor of AGT, $O^6$-BzG 6 h post-transfection. Note that, PBMCs being unattached, can't be visible on the same plane during imaging with that of the MDA-MB-231 cells, which grow on the surface of the petri-dish. Note that, $O^6$-BzG itself was cytotoxic.

**A**

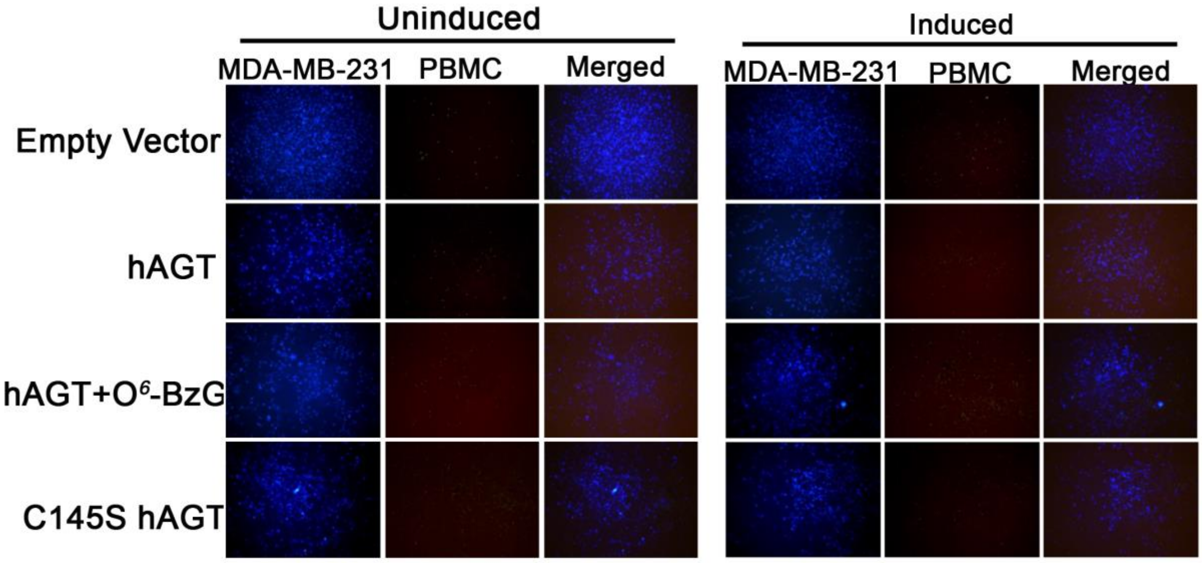


**B**

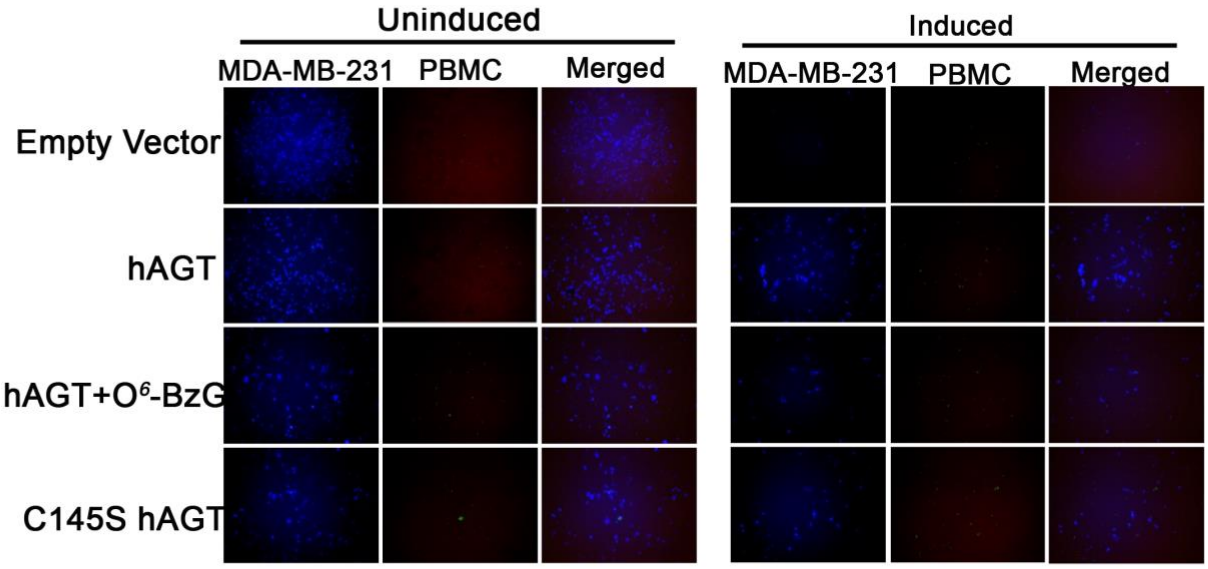

**Figure S3:** Detection of hAGT or C145S-hAGT in the cell lysate of MDA-MB-231 cells transfected with hAGT pEGFP-N1 or C145S hAGT pEGFP-N1.

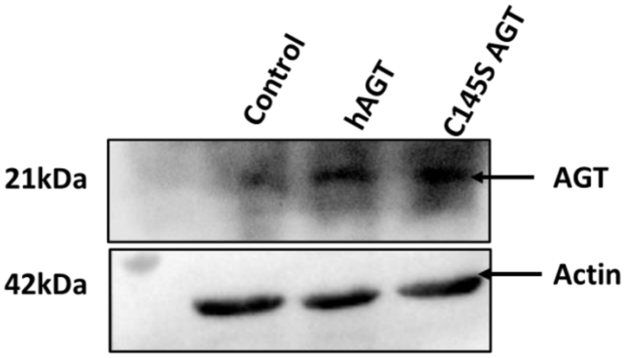

**Figure S4:** Differentiation of THP-1 monocytes to M1 macrophages. A) DIC image of THP-1 cells where control is undifferentitated monocytes, day 3 and 5 are differentiated macrophages after 3 and 5 days of PMA exposure. B) QRT-PCR of CD68 and CD14 markers in THP-1 cells indicating differentiation into macrophages. C) M1 polarisation of macrophages, determined by the increased expression of NOS2 (M1-polarisation marker) and absence of expression of ARG1 (M2-polarisation marker) in western blot. Actin was used as loading control.

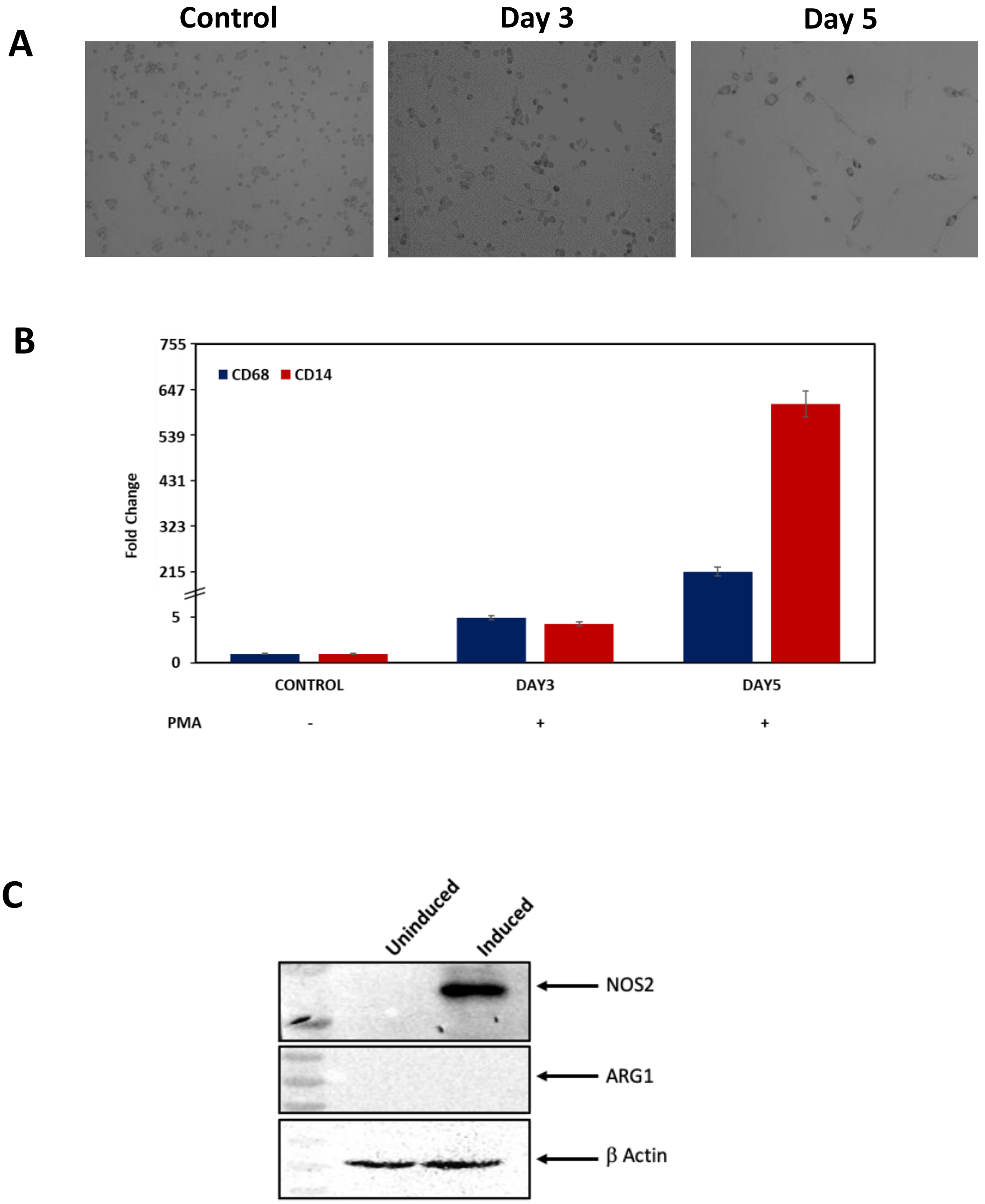

**Figure S5.** Effect of hAGT on induced macrophage-mediated clearance of target epithelial cells. Live cell images of co-cultured MDA-MB-231 and induced macrophage cells that were incubated for A) 0 h and B) 36 h and labelled with cell tracker blue and orange dyes, respectively. MDA-MB-231 cells were transfected with empty pEGFP-N1, hAGT pEGFP-N1 or mutated C145S hAGT pEGFP-N1 vectors for overexpression of the corresponding proteins and subsequently co-cultured with induced macrophage cells. In one set of experiments, hAGT pEGFPN-1-expressing MDA-MB-231 cells were also treated with the known inhibitor of AGT *O*[6]-BzG 6 h post-transfection. Note that, *O*[6]-BzG itself was cytotoxic.

**A**

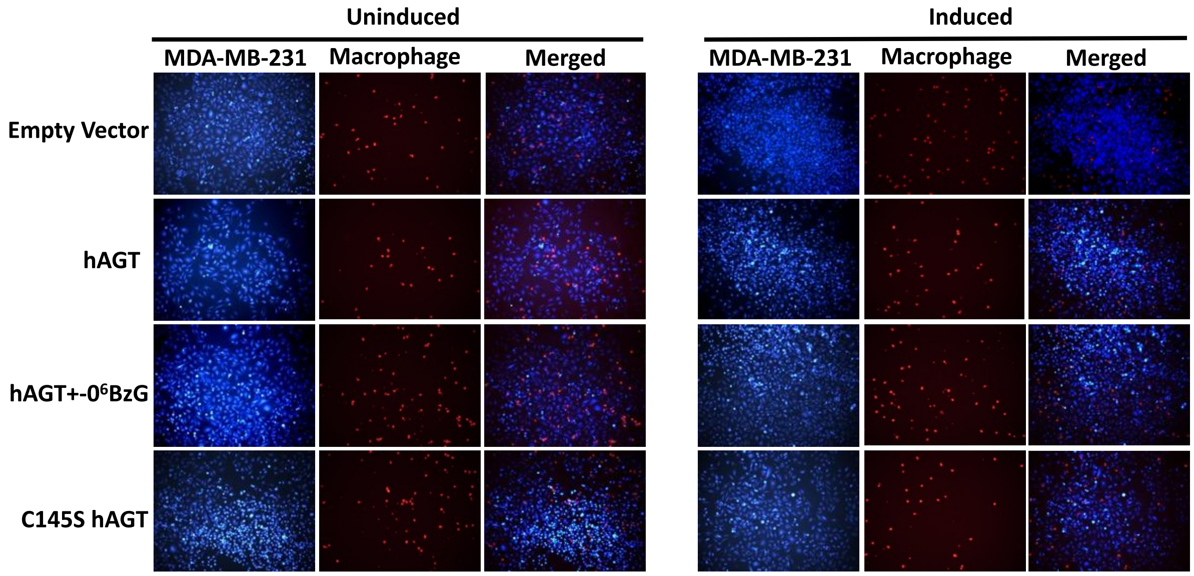


**B**

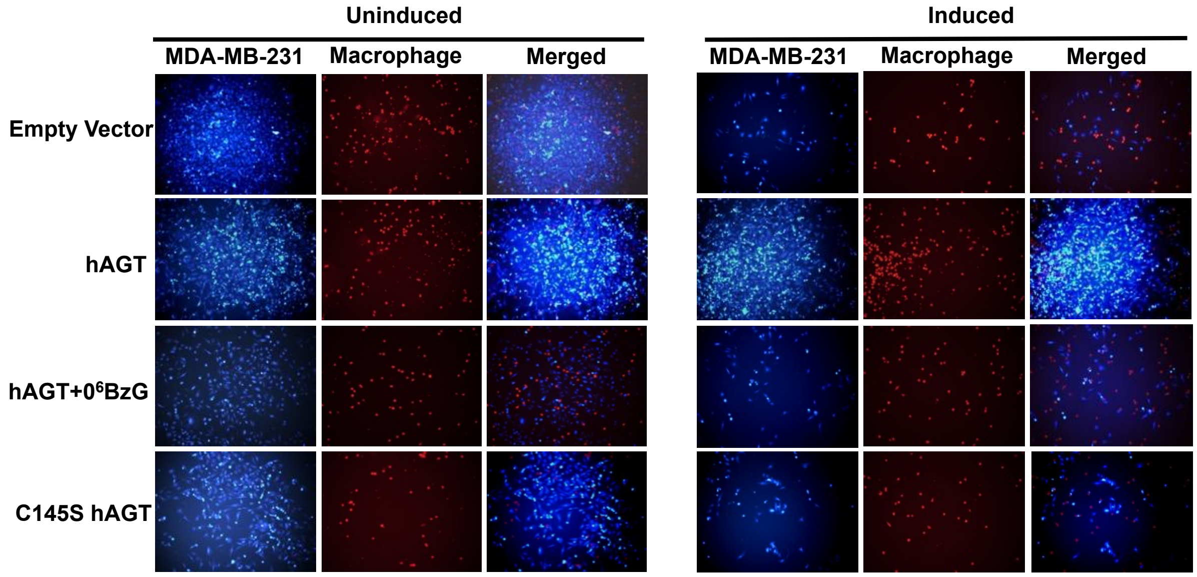

**Figure S6**: Coomassie stain images of recombinant and purified hAGT and C145S hAGT

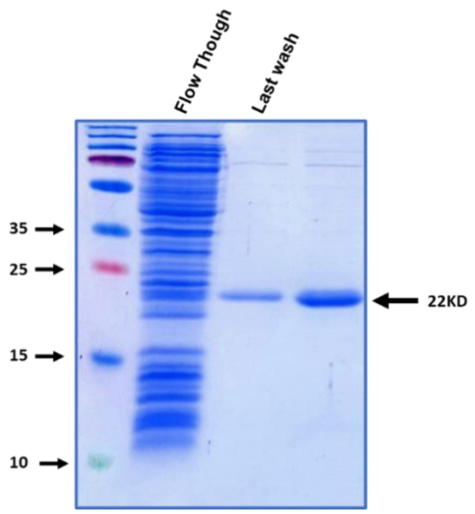
